# Comment on “A genetic signature of the evolution of loss of flight in the Galapagos cormorant”

**DOI:** 10.1101/181826

**Authors:** Mark J. Berger, Gill Bejerano

## Abstract

Burga et al. could not identify gene regulatory regions which might contribute to flightlessness in the Galapagos cormorant. Using a bird-specific alignment we discover 48 limb enhancers showing strong accelerated evolution in *P. harrisi*, including enhancers of key cilia development, hedgehog signaling, and planar cell polarity genes, such as *Prickle1*, *Twist2* and *Ndk1*, extending Burga's proposed mechanism into the non-coding genome.

---

Burga et al. (1) sequenced the genomes of four cormorant species to identify candidate mutations which may have contributed to flightlessness in the Galapagos cormorant. The authors found numerous function-altering variants in genes related to cilia development, hedgehog signaling, and the planar cell polarity (PCP) pathway, such as *Ift122, Gli2,* and *Fat1.* They also identified a 4-amino acid deletion in *Cux1,* a highly-conserved homeodomain transcription factor. The authors demonstrate that *Cux1* regulates the expression of key cilia and PCP genes, as well as that the 4-amino acid deletion impairs *Cux1* from activating gene expression. These results suggest that perturbations of cilia development, hedgehog signaling, and the PCP pathway may play a role in reduced keel and wings of the Galapagos cormorant.

While the authors implicated many protein-coding mutations in the disruption of the primary cilium, they found no evidence of a non-coding signature. The authors aligned phastCons (2) conserved regions from the UCSC 100-way vertebrate multiple alignment (3) to each of the four cormorant genomes using reciprocal best BLAST alignments. Using PhyloP (4), the authors tested each of the resulting 40,812 candidate regions for accelerated evolution in *P. harrisi* compared to all tetrapods. With a false-discovery rate (FDR) cutoff of 5%, Burga et al. found only 11 accelerated regions in *P. harrisi,* none of which overlapped limb enhancer marks.

Here, we show that by testing regions which are conserved among avian species, instead of those conserved among vertebrates, we discover 48 accelerated non-coding regions which overlap limb enhancer marks. Using the four cormorant species sequenced by the authors, as well as *Phalacrocorax carbo, Nipponia nippon,* and *Gallus gallus* galGal5 assembly as reference (Figure 1A), we assembled a genome-wide multiple alignment between the seven species using MULTIZ (5). A neutral model of evolution was constructed using fourfold degenerate sites in chicken derived from the Ensembl 89 gene set (6). Ignoring *P. harrisi,* phastCons (2) identified 894,840 highly conserved regions not overlapping a protein-coding gene in chicken. Non-coding conserved regions within 100bp of each other were merged, and regions smaller than 100bp were removed, resulting in 430,015 candidate regions (Figure 1B).

**Figure 1.**
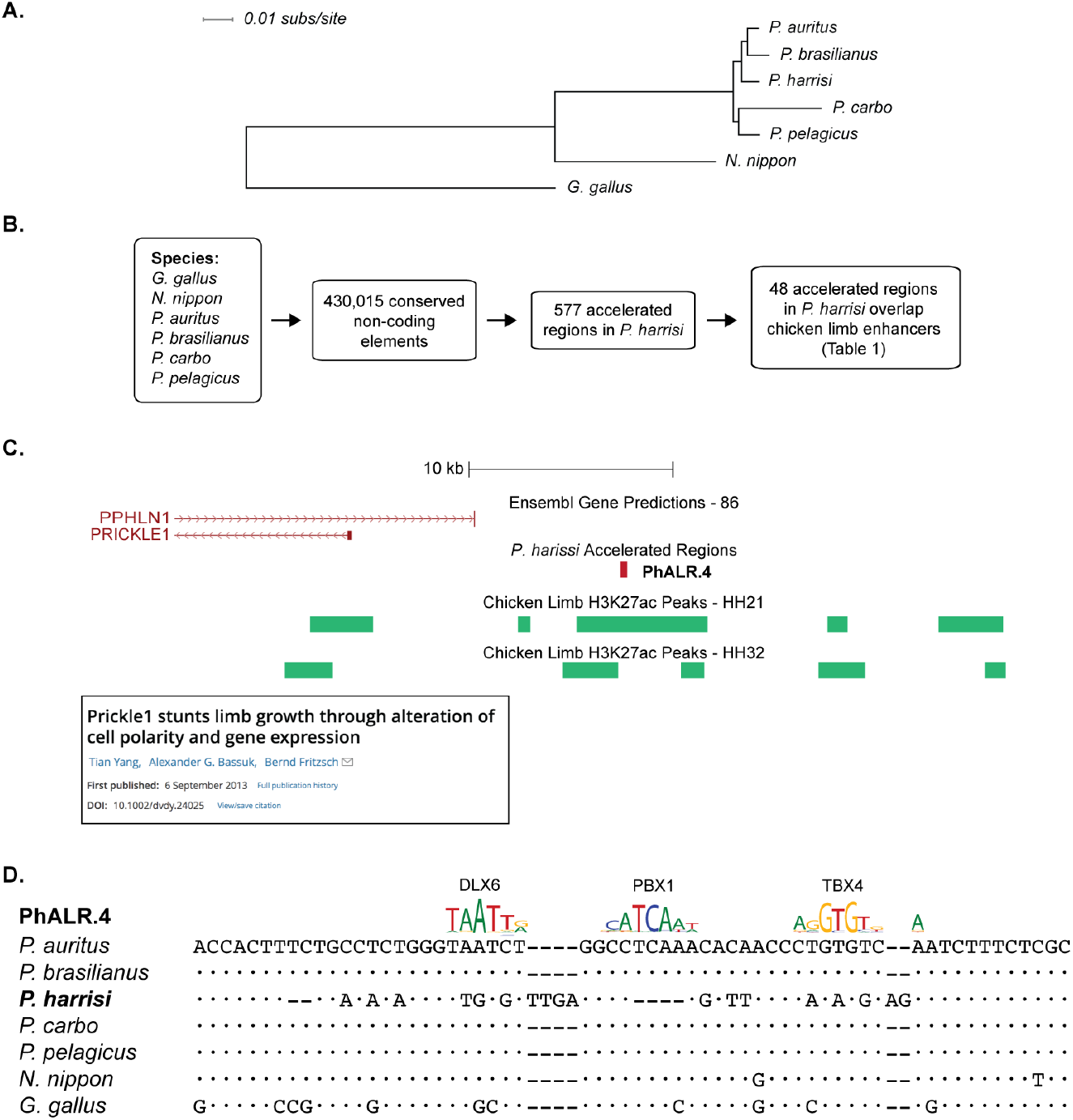
Identifying accelerated limb enhancers in the Galapagos cormorant. (A) Phylogeny of our bird-specific multiple alignment. (B) Ignoring *P. harrisi,* we identified 430,015 conserved non-coding elements. Of these, 577 elements are significantly accelerated in the Galapagos cormorant. 48 of these overlap chicken limb enhancer marks. (C) For example, PhALR.4 lies 13kb upstream of *PRICKLE1.* Mouse *Prickle1* mutants experience stunted limb growth (8) and primary cilia defects (9). (D) A portion of PhALR.4 is shown highly diverged in *P. harrisi.* Bases which are identical to the sequence in *P. auritus* are represented by a dot, the absence of a basepair is denoted by a dash. Despite the very short evolutionary distance between *P. auritus*and *P. harrisi* (Figure 1A), the corresponding sequence in *P. harrisi* is even more diverged than the corresponding sequence in *G. gallus.* Using position weight matrices (15), we predicted three binding sites of important limb development regulators which appear eroded in the Galapagos cormorant.

Using PhyloP (4) we asked whether any of these candidate regions have undergone accelerated molecular evolution in the Galapagos cormorant. Even with a stringent FDR cutoff of 1%, we found 577 accelerated regions in *P. harrisi* (Figure 1B). Of these, 48 elements overlap H3K27Ac enhancer marks at chicken developmental time points HH21 or HH32 by at least 50% (Table 1) (7). We refer to this set of 48 regions as *P. harrisi* Accelerated Limb Regions (PhALRs).

**Table 1.** *P. harrisi* Accelerated Limb Regions (PhALRs). Gene regulatory domains are assigned using the default rules of Genomic Regions Enrichment of Annotations Tool (GREAT) v3.0.0. The nearest transcription start sites are reported on either side of each element. Negative distances are upstream of each transcription start site and positive distances are downstream.

| galGal5 Chrom | Start | End | Element Name | Nearest genes (Distance to TSS) |
| --- | --- | --- | --- | --- |
| chr1 | 1174874 | 1175514 | PhALR.1 | TUBAL3 (-23.3k), NET1 (-26.5k) |
| chr1 | 6994904 | 6995093 | PhALR.2 | FRMD4A (-90.4k), ENSGALG00000033226 (+357.5k) |
| chr1 | 25813481 | 25813762 | PhALR.3 | MDFIC (-555) |
| chr1 | 29721579 | 29721941 | PhALR.4 | PRICKLE1 (-13.1k), ADAMTS20 (+286.2k) |
| chr1 | 34364240 | 34364404 | PhALR.5 | LLPH (+4.9k), HMGI-C (+160.3k) |
| chr1 | 35924280 | 35924467 | PhALR.6 | ENSGALG00000040855 (+20.2k), PTPRB (-32.5k) |
| chr1 | 57866756 | 57866951 | PhALR.7 | CHRM2 (-1.1k) |
| chr1 | 62063564 | 62064074 | PhALR.8 | ENSGALG00000013042 (-24.9k), PEX26 (-50.5k) |
| chr1 | 104823969 | 104824070 | PhALR.9 | MIS18A (+57.3k), HUNK (+91.9k) |
| chr10 | 18730427 | 18730944 | PhALR.10 | SMAD6 (+31.2k), SMAD3 (-81.7k) |
| chr11 | 6499531 | 6499950 | PhALR.11 | NKD1 (-31.1k), BRD7 (-130.7k) |
| chr11 | 12075244 | 12075577 | PhALR.12 | CDH11 (-257.7k), CDH5(-372.0k) |
| chr11 | 17481889 | 17482561 | PhALR.13 | ENSGALG00000029278 (+102.0k),<br>ENSGALG00000039008 (+209.6k) |
| chr12 | 9433627 | 9433759 | PhALR.14 | EEFSEC (+80.4k), ENSGALG00000044250 (-129.2k) |
| chr13 | 13174020 | 13174369 | PhALR.15 | TCOF1 (-5.4k), ARSI (-9.6k) |
| chr14 | 4240133 | 4240298 | PhALR.16 | RNF216 (+30.5k), FSCN1 (+33.5k) |
| chr15 | 1350352 | 1350509 | PhALR.17 | ZDHHC8 (+24.5k), PGAM5(-71.6k) |
| chr15 | 12355200 | 12355870 | PhALR.18 | TBX5 (-25.2k), TBX3 (+77.6k) |
| chr2 | 13968283 | 13969054 | PhALR.19 | ITGB1 (-77.5k), NRP1(+82.1k) |
| chr2 | 20451532 | 20451702 | PhALR.20 | FAM171A1 (-68.8k), ITGA8 (+87.4k) |
| chr2 | 24132193 | 24133200 | PhALR.21 | SLC25A13 (+110.9k), DYNC1I1 (+146.9k) |
| chr2 | 35320405 | 35320605 | PhALR.22 | SATB1 (+13.9k), TBC1D5 (-293.3k) |
| chr2 | 61118995 | 61119310 | PhALR.23 | JARID2 (-146.7k), CD83(+226.0k) |
| chr2 | 62397629 | 62398027 | PhALR.24 | TBC1D7 (-18.5k), GFOD1(+60.8k) |
| chr2 | 64221316 | 64222213 | PhALR.25 | SLC35B3 (-116.7k), ENSGALG00000042845 (+438.6k) |
| chr2 | 86816902 | 86817718 | PhALR.26 | IRX2 (+90.1k), IRX4(-485.9k) |
| chr21 | 3120789 | 3121217 | PhALR.27 | ENSGALG00000030881 (+27.1k),<br>ENSGALG00000002232 (-79.0k) |
| chr23 | 2021824 | 2022005 | PhALR.28 | AHDC1 (+5.2k), WASF2 (-25.0k) |
| chr23 | 5507699 | 5507877 | PhALR.29 | HEYL (-6.9k), NT5C1A (+10.7k) |
| chr26 | 4264439 | 4264822 | PhALR.30 | ANKS1A (+7.3k), ENSGALG00000002692 (+87.0k) |
| chr28 | 4774298 | 4774715 | PhALR.31 | KDM4B (+68.6k), PTPRS(+148.4k) |
| chr3 | 14230323 | 14230448 | PhALR.32 | PLCB4(-183.5k), PLCB1(+349.7k) |
| chr3 | 54386692 | 54387333 | PhALR.33 | NHSL1 (+12.5k), ENSGALG00000035680 (+178.5k) |
| chr3 | 56803537 | 56804397 | PhALR.34 | EYA4 (-78.3k), RPS12(+161.6k) |
| chr33 | 269239 | 269347 | PhALR.35 | CERS5 (-5.0k), LIMA1 (+17.3k) |
| chr4 | 32409496 | 32409661 | PhALR.36 | NR3C2 (+161.5k), ARHGAP10 (+169.1k) |
| chr4 | 32486613 | 32486885 | PhALR.37 | NR3C2 (+84.3k), ARHGAP10(+246.3k) |
| chr4 | 39319472 | 39319936 | PhALR.38 | TACR3 (-129.2k), TET2 (-212.5k) |
| chr4 | 66532631 | 66532867 | PhALR.39 | SPATA18 (+54.3k), USP46 (+119.3k) |
| chr5 | 28082372 | 28082505 | PhALR.40 | GALNT16 (+19.7k), ERH(+56.5k) |
| chr6 | 12739800 | 12740116 | PhALR.41 | ZMIZ1 (-327.4k), RPS24(+418.0k) |
| chr6 | 33850891 | 33851286 | PhALR.42 | MGMT (-178.5k), ENSGALG00000025996 (+443.9k) |
| chr7 | 4535095 | 4535543 | PhALR.43 | MREG (+28.8k), FN1(-95.9k) |
| chr7 | 6116197 | 6116726 | PhALR.44 | TWIST2 (-11.0k), ASB1 (+142.6k) |
| chr7 | 33589086 | 33589506 | PhALR.45 | Sip1 (-892) |
| chr8 | 15800494 | 15800807 | PhALR.46 | LMO4 (+864) |
| chr9 | 20089307 | 20089868 | PhALR.47 | MECOM (+26.6k), ENSGALG00000031382 (+69.3k) |
| chrZ | 79330495 | 79331788 | PhALR.48 | ZNF608 (-293.3k), GRAMD3 (-345.5k) |

A number of the resulting regions corroborate Burga et al.’s purposed model. For example, PhALR.4 lies 13kb upstream of *Prickle1* (Figure 1C) and is highly diverged in the Galapagos cormorant (Figure 1D). *Prickle1* is a key member of the PCP pathway, and mice mutants with a premature stop codon in the 3^rd^ LIM domain exhibit defective limb growth (8). Mutants also exhibit altered expression of key developmental limb genes, such as *Bmp4* and *Wnt5a* (8). Furthermore, mice with a missense mutation in *Prickle1* display primary cilium defects (9). PhALR.11 lies 31kb upstream of *Ndk1,* a highly-conserved gene which regulates Wnt signaling. Knockdown of *nkd1* in zebrafish leads to loss of cilia in Kupffer’s vesicle (10).

Other regions suggest that additional mechanisms contribute to the reduced keel and wings in the Galapagos cormorant. PhALR.44 lies 11kb upstream of *Twist2. Twist2* is a bHLH transcription factor which represses *Grem1* to terminate the Shh/Grem1/Fgf autoregulatory loop in early limb development (11). Viral-induced misexpression of *Twist2* results in underdeveloped limbs in mice (11). PhALR.21 lies in the intron of *Dync1i1,* proximal to exon 15. Exon 15 and 17 of *DYNC1I1* have been shown to function as distal enhancers for *DLX5* and *DLX6,* two critical regulators of limb development (12). Chromosomal abnormalities of *DYNC1I1,* as well as simultaneous disruption of *Dlx5* and *Dlx6,* both lead to split hand and foot malformation in humans and mice, respectively (12, 13). Other *P. harrisi* Accelerated Limb Regions are found next to *SMAD, TBX, IRX* genes and more (Table 1).

Burga et al. identified genes in *P. harrisi* with striking loss of function mutations and demonstrated that cilia development, hedgehog signaling, and the PCP pathway may be responsible for flightlessness in the Galapagos cormorant (1). Developmental genes are highly pleiotropic and will only accumulate deleterious mutations when the organism can withstand the corresponding loss in fitness throughout all developmental contexts (14). It is plausible that additional genes contribute to flightlessness in the Galapagos cormorant, but present with no loss of function coding mutations. Instead, the oft less pleiotropic cis-regulatory elements for these genes may mutate more freely. Here, we discover multiple putative limb enhancers which have diverged significantly in the Galapagos cormorant, including those upstream of functionally relevant genes. Our results corroborate the model purposed by Burga et al., and extend it into the non-coding genome.

